# Protocol for standardized minimally invasive mouse models of bisphosphonate-related and radiation-induced jaw osteonecrosis

**DOI:** 10.64898/2026.06.28.735116

**Authors:** Zhangfan Ding, Jiayuan Zhang, Hanyu Liu, Abhishek Chandra, Makarand V Risbud, Anjali P Kusumbe, Junyu Chen

**Affiliations:** State Key Laboratory of Oral Diseases, National Center for Stomatology, National Clinical Research Center for Oral Diseases, West China Hospital of Stomatology, Sichuan University, Chengdu 610041, China; 2Tissue and Tumor Microenvironments Lab, Cancer Discovery and Regenerative Medicine Program, Lee Kong Chian School of Medicine, Nanyang Technological University, 636921 Singapore; Multidisciplinary Institute of Ageing (MIA-Portugal), University of Coimbra, Coimbra 3004-504, Portugal; Department of Physiology and Biomedical Engineering, Mayo Clinic, Rochester, MN 55905, USA; Robert and Arlene Kogod Center on Aging, Mayo Clinic, Rochester, MN 55905, USA; Department of Biochemistry and Molecular Biology, Mayo Clinic, Rochester, MN 55905, USA; Department of Orthopaedic Surgery, Sidney Kimmel Medical College, Thomas Jefferson University, Philadelphia, PA, USA; Graduate Program in Cell Biology and Regenerative Medicine, Thomas Jefferson University, Philadelphia, PA, USA

## Abstract

This protocol describes a standardized and reproducible minimally invasive approach for establishing mouse models of bisphosphonate-related osteonecrosis of the jaw (BRONJ) and osteoradionecrosis of the jaw (ORNJ). The method combines a unified low-trauma oral surgical procedure with disease-specific injury induction strategies to generate robust and clinically relevant models of jaw osteonecrosis. For BRONJ, systemic zoledronic acid administration is coupled with mandibular first molar extraction using tape-assisted mouth opening and customized bent micro-forceps, minimizing soft tissue damage and reducing procedural variability. For ORNJ, a customized lead-shielding platform enables precise, noninvasive mandible-targeted irradiation, producing reproducible bone injury while limiting off-target radiation exposure. Together, these complementary models provide a consistent and minimally invasive framework for investigating jaw osteonecrosis arising from distinct etiologies. The protocol supports comprehensive downstream analyses, including micro-computed tomography, histology, and immunofluorescence, and facilitates mechanistic studies of disease pathogenesis, bone regeneration, and therapeutic intervention.

## BEFORE YOU BEGIN

Osteonecrosis of the jaw (ONJ) represents a severe and debilitating complication associated with widely used clinical therapies. It primarily includes two major clinical subtypes: bisphosphonate-related osteonecrosis of the jaw (BRONJ) and osteoradionecrosis of the jaw (ORNJ).

Bisphosphonates are widely used in the treatment of skeletal disorders, including osteoporosis and multiple myeloma ^1,2^, while radiotherapy remains a cornerstone in the management of head and neck malignancies ^3–5^. Despite their clinical benefits, both therapies are associated with severe late complications in the maxillofacial region, collectively manifesting as jaw osteonecrosis. Bisphosphonate-related osteonecrosis of the jaw (BRONJ) and osteoradionecrosis of the jaw (ORNJ) are clinically distinct but pathophysiologically related conditions characterized by progressive bone destruction, chronic infection, fistula formation, and pathological fracture, which severely impair oral function and quality of life ^3,6^.

Although these conditions are increasingly recognized, their underlying mechanisms remain incompletely understood, and effective disease-modifying therapies are still lacking. A major limitation in the field is the absence of standardized and reproducible preclinical models that reliably recapitulate key pathological features across different etiologies. Existing experimental approaches are often heterogeneous and technically demanding, leading to substantial variability and limited cross-study comparability ^4,7–9^.

To address this gap, we describe standardized experimental protocols for establishing minimally invasive mouse models of BRONJ and osteoradionecrosis of the jaw (ORNJ). These protocols are designed to improve procedural consistency and reproducibility while reducing technical variability. The models support downstream applications including micro-CT imaging, histological assessment, and molecular analyses, and provide a robust experimental framework for studying the mechanisms and therapeutic interventions of jaw osteonecrosis.

### Institutional permissions

All the animals utilized for the studies were housed at Sichuan University. All animal procedures were reviewed and approved by the Research Ethics Committee of West China Hospital of Stomatology (Approval Nos. WCHSIRB-D-2023-245, WCHSIRB-D-2024-248). Adult male C57BL/6 mice were obtained from the Chengdu Dashuo Bio-Technology Co., Ltd. (China).

### Innovation

This protocol establishes a standardized and reproducible minimally invasive approach for generating mouse models of bisphosphonate-related osteonecrosis of the jaw (BRONJ) and osteoradionecrosis of the jaw (ORNJ), improving procedural consistency and reducing inter-operator variability. A unified oral surgical strategy is applied across both models, integrating tape-assisted mouth opening and blunt tissue retraction to enable stable and atraumatic exposure of the mandibular molar region. In BRONJ, systemic zoledronic acid administration is combined with controlled mandibular first molar extraction using bent micro-forceps to achieve precise and reproducible tooth removal. In ORNJ, a customized lead shielding system enables noninvasive, mandible-targeted irradiation with improved spatial precision and reduced off-target effects. Together, these refinements provide a robust and reproducible framework for downstream micro-CT, histological, and immunofluorescence analyses and for mechanistic studies of jaw osteonecrosis and therapeutic evaluation.

## KEY RESOURCES TABLE

| REAGENT or RESOURCE | SOURCE | IDENTIFIER |
| --- | --- | --- |
| **Chemicals, peptides, and recombinant proteins** |  |  |
| Zoledronic acid (ZA) | APExBIO | A1352 |
| NaCl | Sigma | S7653 |
| Tribromoethyl alcohol (Avertin) | Biotopped | Top9153S |
| Iodine (povidone-iodine) | Sigma-Aldrich | PVP-I |
| Ethanol | Sigma-Aldrich | 1070172511 |
| PBS (10×) | VWR | 437117K |
| Donkey serum | Solarbio | SL050 |
| Paraformaldehyde (PFA) | Sigma-Aldrich | P6148 |
| EDTA | Sigma-Aldrich | E6758 |
| Polyvinylpyrrolidone | Sigma-Aldrich | P5288 |
| Sucrose | Sigma-Aldrich | V900116 |
| Gelatin | Sigma-Aldrich | G1890 |
| Fluoromount-G | Invitrogen | 00-4958-02 |
| Triton X-100 | Sigma-Aldrich | T8787 |
| H&E staining kit | Servicebio | G1005-100ML |
| DAPI | Sigma-Aldrich | D9542 |
| **Antibodies** |  |  |
| CD31 | Abcam | ab24590 |
| Endomucin | Santa Cruz<br>Biotechnology | sc-65495 |
| Sp7 | Abcam | ab209484 |
| Donkey anti-Mouse IgG (H+L) Alexa Fluor 488 | Invitrogen | A-21202 |
| Donkey anti-Rat IgG (H+L) Alexa Fluor Plus 555 | Invitrogen | A48270 |
| Donkey anti-Rabbit IgG (H+L) Alexa Fluor 647 | Invitrogen | A-31573 |
| **Equipment** |  |  |
| Ophthalmic scissors (straight, 10 cm) | Biosharp | BS-SOR-S-100 |
| Micro-forceps (straight, 14 cm) | Biosharp | BS-ZER-S-014A |
| Blunt tissue forceps | Biosharp | BS-ZER-S-100 |
| Animal bowl | Servicebio | QX0030 |
| Low-profile blades | Leica | 14035838382 |
| Cover glass | VWR | 631-0135 |
| Micro-CT scanner | SCANCO | μCT45 |
| Cryostat | Leica | CM1950 |
| Digital slide scanner | Olympus | SLIDEVIEW<br>VS200 |
| Spinning disk confocal microscope | Olympus | SpinSR |
| Heating pad | N/A | N/A |
| **Software** |  |  |
| Dataviewer (version 1.5.6.2) | Bruker | N/A |
| CTVox (version 1.18.4.0) | Bruker | N/A |
| OlyVIA (version 4.1) | Olympus / Evident<br>Scientific | N/A |
| Imaris (version 10.1.0) | Bitplane | N/A |
| Imaris File converter (version 10.1.0) | Bitplane | N/A |
| Adobe Illustrator CC | Adobe | N/A |

**CRITICAL:** PFA is toxic. The fixative solution should be prepared inside a fume hood.

## Materials and equipment

### Injection solution

Dissolve ZA powder in 8 mL 0.9% NaCl solution (normal saline).

**CRITICAL:** This solution should be stored at 4°C for a maximum of 1 month.

**Fixation solution**

| REAGENT | FINAL CONCENTRATION | AMOUNT |
| --- | --- | --- |
| Paraformaldehyde | 4% | 4 g |
| PBS (1×) | 1× | 100 mL |

Store at 4 °C for up to 1 weeks. Prepare fresh for best results.

**CRITICAL:** PFA is highly toxic and should be manipulated in the fume hood, and the appropriate PPE is required.

### Decalcifying solution (10% EDTA, pH 7.4)

1. Dissolve 10 g EDTA in 80 mL distilled water, adjust pH to 7.4 with NaOH, then bring to 100 mL.
2. Store at room temperature for up to 1 month.

**Antibody solution**

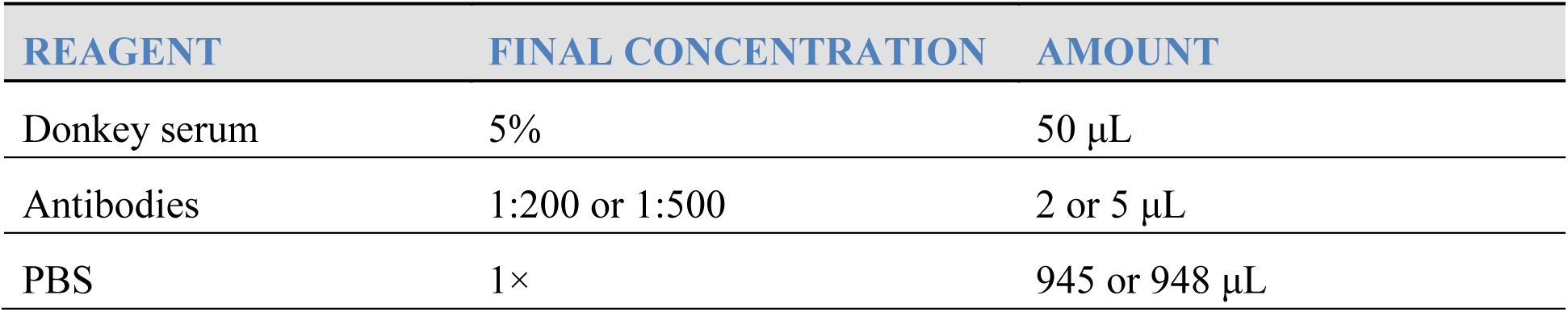

**Antibody dilutions**

| ANTIBODY | DILUTION |
| --- | --- |
| CD31 | 1:200 |
| Endomucin | 1:200 |
| Sp7 | 1:200 |
| Donkey anti-Mouse IgG Alexa Fluor 488 | 1:500 |
| Donkey anti-Goat IgG Alexa Fluor 555 | 1:500 |
| Donkey anti-Rabbit IgG Alexa Fluor 647 | 1:500 |

## STEP-BY-STEP METHOD DETAILS

### Preoperative management

This section describes the preoperative procedures for BRONJ and ORNJ, followed by tooth extraction, postoperative care, and downstream analyses. Preoperative management is divided into Part 1 for BRONJ and Part 2 for ORNJ.

### Part 1. For BRONJ

Preoperative drug injection

### Timing: [6 weeks]

This section describes the process of drug injection into mice for the model of mandibular bone necrosis.

1. House mice (8-12 weeks old male C57BL/6J) in standard cages with a 12 h light/dark cycle and ad libitum access to food and water.
2. Weighing each mouse. The mice are injected intraperitoneally with the prepared solution at a dose of 200 µg/kg of ZA, three times a week.

**Note:** Each injection is performed using a new disposable needle. The tooth extraction surgery schedule to be carried out at the weekend of the 6^th^ week after the first drug injection.

### Part 2. For ORNJ

S1. Mouse anesthesia

### Timing: [10 minutes]

This section describes the process of animal anesthesia.

3. House mice (8-12 weeks old male C57BL/6J) in standard cages with a 12 h

light/dark cycle and ad libitum access to food and water.

4. Weighing each mouse. The mice are injected intraperitoneally with the prepared solution at a dose of 0.2 mL/10g of tribromoethyl alcohol (Avertin).

S2. Mouse positioning

### Timing: [5 minutes]

This section describes how to fix the mouse in a customized lead chamber, which enables precise radiation exposure of mandible.

5. Place the mouse in the customized lead chamber in a supine position (Figure 1A). Close the lid and leave a square opening to expose only the mandible. Adjust the area of the gap according to the size of the mandible (Figure 1B-D). The chamber can accommodate 4 mice at a time.

**Figure 1.**
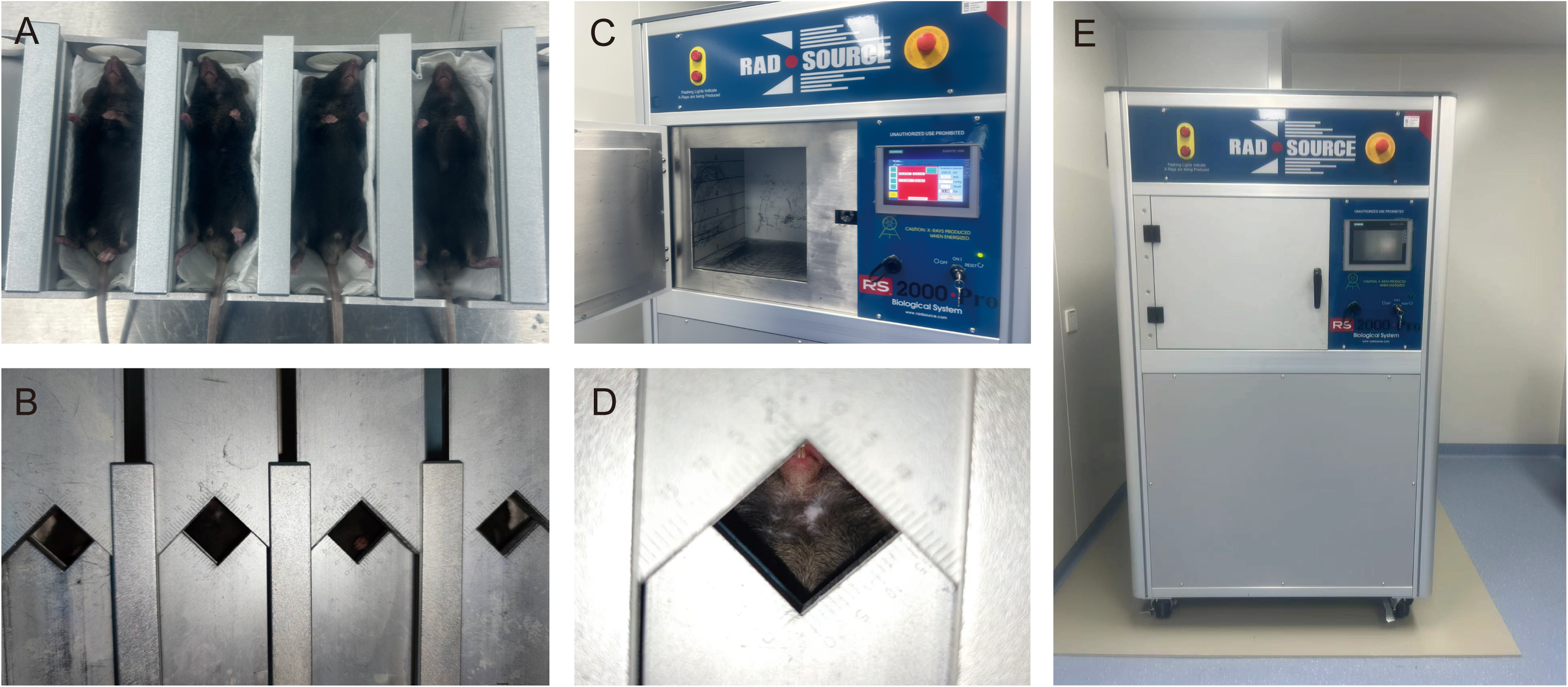
The establishment and validation of the mouse ORNJ model. (A) Placement of the mouse in the customized lead chamber. (B) Close the lid to protect the mice from excess radiation injury. (C) The radiation cabin which accommodates four mice. (D) The gap on the lid for exposing the area of mouse mandible. (E) Set the radiation dose and start the irradiator.

S3. Radiation treatment

### Timing: [6 minutes]

This section describes how to administer radiation exposure to the mice.

6. Set the single radiation dose to 16 Gy and start radiation (Figure 1E). Take out the mice from the chamber and transfer them to the cage.

**Note:** Leave the room where the X-ray irradiator is located to avoid the impacts of radiation on operators.

S4. Post-radiation care

**Timing: [2 weeks]**

This section describes the managements after radiation.

7. Feed the mice in standard cages with a 12 h light/dark cycle and ad libitum access

to food and water.

Surgical instrument preparation

**Timing: [20 minutes] (excluding the time for sterilization)**

This section describes how to prepare the necessary equipment for an extraction procedure, which is carried out at the end of the 6th week of drug injection.

8. Before the tooth extraction surgery, prepare the following instruments in advance and place them in the high-pressure sterilization and disinfection surgical instruments:

A. micro-forceps, the tip of which should be bent beforehand (Figure 2B), to grasp the bifurcation when extracting the mandibular first molar (M1) and loosen the tooth before extraction.
B. Blunt tissue forceps, one of which used as a buccal retractor when exposing the mandibular molar area and another used as a mouth opener (Figure 2D).
C. Disposable syringe needles (at least 2 per mouse) to fix the mouth opener and separate the gingiva.
D. Sterile cotton balls used for stopping bleeding and cleaning the surgical site (Figure 2A).
E. Ophthalmic scissors to cut tape.

**Figure 2.**
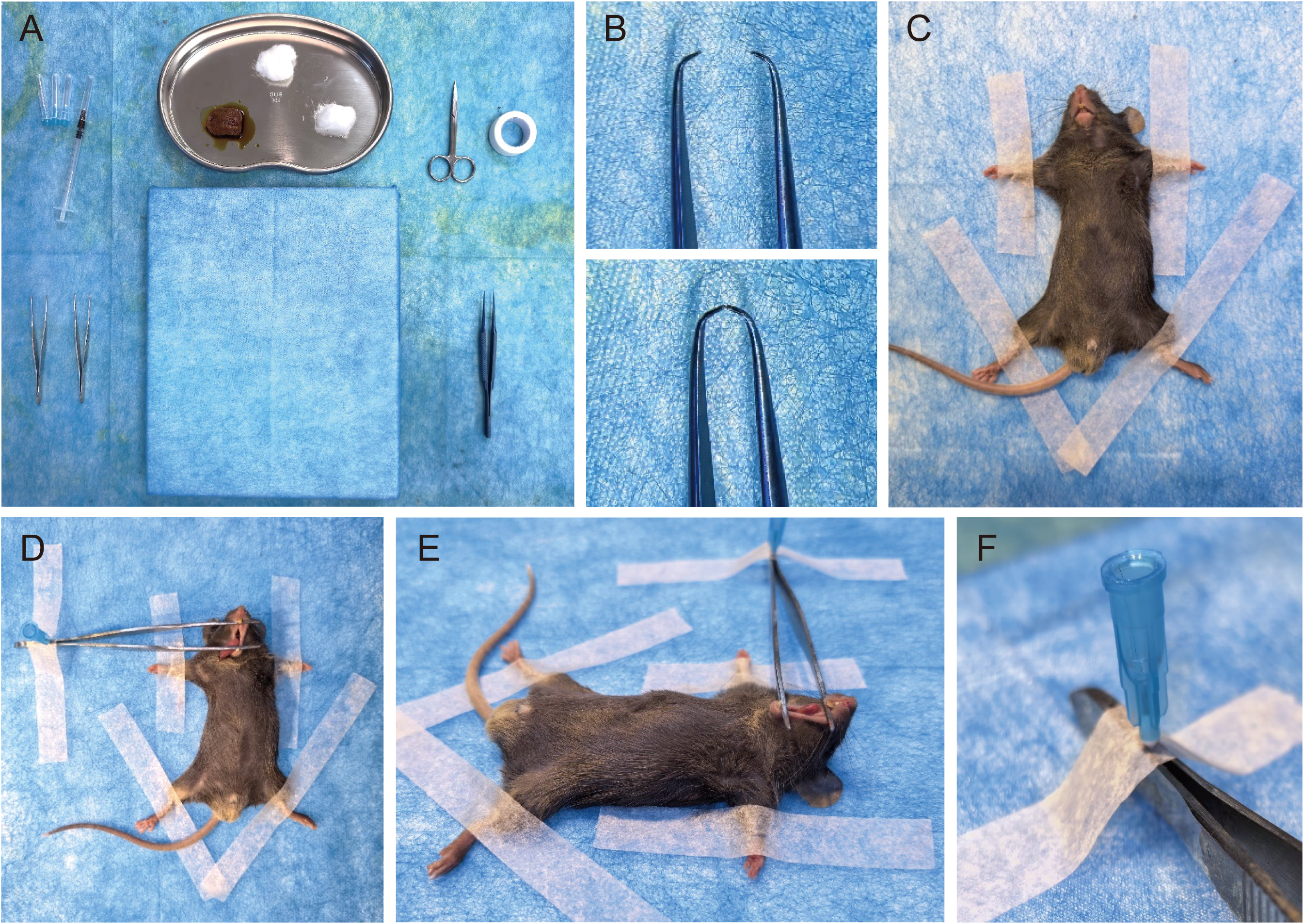
The surgical preparation and positioning of the mouse before tooth extraction. (A) Complete surgical platform and required instruments. (B) The tip of the bent micro-forceps. (C-F) Methods for immobilizing mice and opening the mouse’s mouth.

**Note:** Use a high-pressure sterilizer to perform a 15-minute sterilization process at a temperature of 121°C to disinfect all surgical instruments and materials.

9. Use disposable surgical drapes to set up the surgical table for mice. Disinfect the surgical table (a foam board, 32×22×2.5cm), injection syringe, tape and heating pad (37 °C, 18×10 cm) with 75% ethanol.

**Note:** Disinfection tape is used to fix the limbs of the mice and the open forceps. A disinfection heating pad is used to maintain the body temperature of the mice during intraoperative and postoperative recovery phases.

10. Disinfect the surgical table with 75% ethanol. Put on the sterile disposable surgical gown, gloves, mask, and head cover. Place the sterilized instruments and materials that have been wiped with 75% ethanol on the disinfected surgical table. **Note:** Before starting the aseptic procedure, make sure that the lighting in the surgical area is sufficient. Using a small table lamp or a head-mounted lamp is usually the common method. Tear sterile cotton balls into small pieces with a diameter of about 5 millimeters for better use.

Animal preparation

**Timing: [10-15 minutes per mouse]**

This section describes how to administer anesthesia to the mice, how to fix the mice and open their mouths to expose the surgical area.

11. Weighing the mice.
12. Anesthetize the mice using 0.2ml/10g Avertin intraperitoneally.

**Note:** Fix the mouse with its abdomen facing upwards. Insert the needle about 1/3 away from the abdominal white line. After the needle tip penetrates the skin, tilt it slightly and insert it into the abdominal cavity and slowly inject the drug. Confirm adequate anesthesia by the absence of pedal reflex (toe-pinch). During the entire surgical procedure, monitor the depth of anesthesia continuously to ensure no reflex reactions occur, especially when extracting a tooth. Maintain body temperature with the heating pad throughout the procedure.

13. Place the mouse on the surgical table and position it in a supine position. Secure the limbs with tape. Use one of the tissue forceps to open the mouth through the upper and lower incisors, and fix the tissue forceps to the surgical table by the disposable syringe needles and the tape (Figure 2C-F).

**Note:** Ensure that the line connecting the upper and lower incisors is parallel to the horizontal plane. The head of the mouse should be directed to the right side so as to facilitate the extraction of the lower molars.

Tooth extraction

**Timing: [5-10 minutes per mouse]**

This section describes how to use the homemade micro-forceps to remove the mandibular M1 from a mouse.

14. Use blunt-tipped forceps to gently open the mouth and expose the area of the left lower molars (Figure 3A-B).
15. Use sterile cotton balls with normal saline to clean the tooth surfaces and gum grooves, to remove debris from the mouth. Use sterile cotton balls with povidone-iodine solution to gently wipe the left M1 area for about 5-10 seconds.

**Figure 3.**
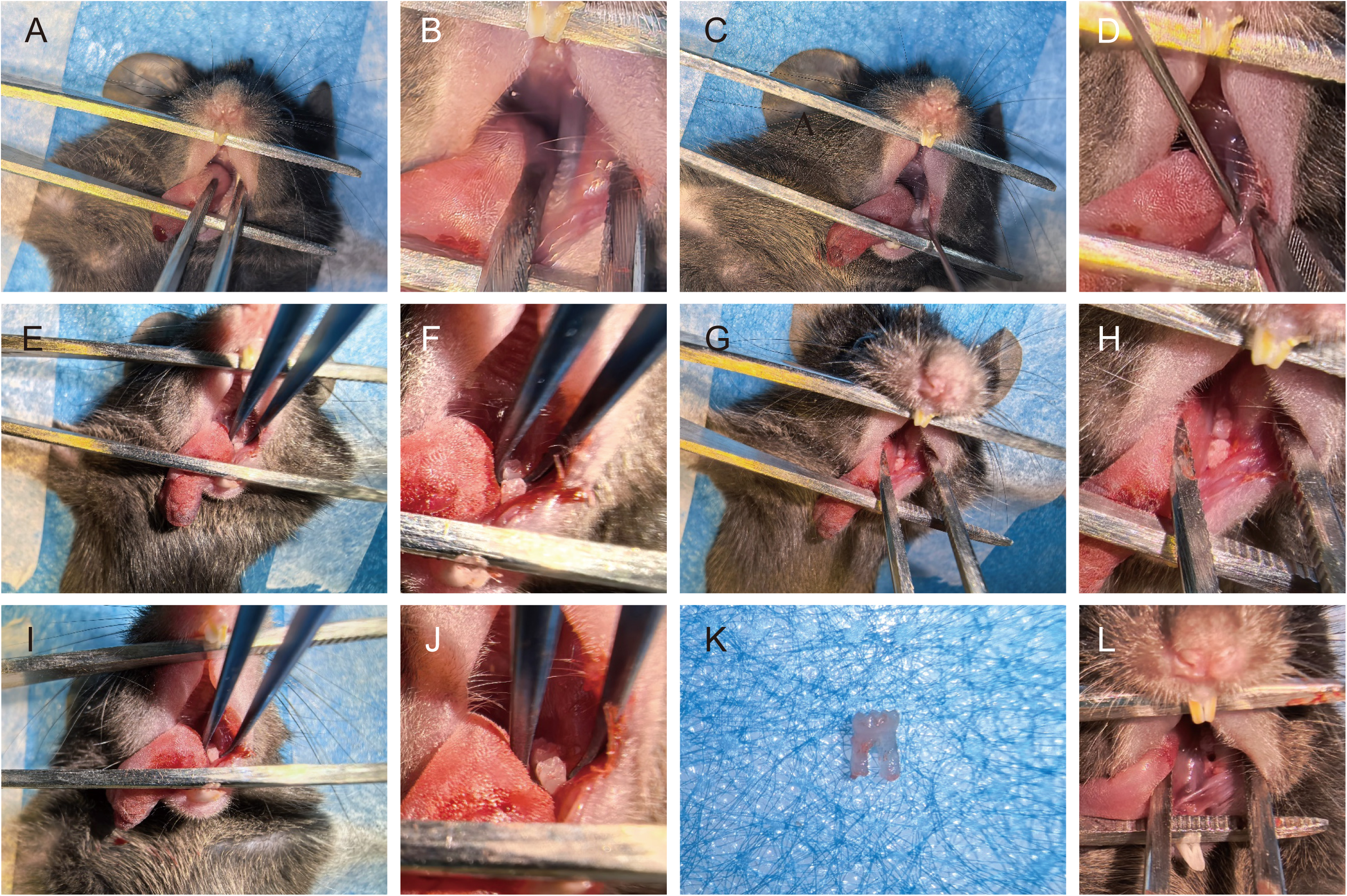
The detailed steps of tooth extraction. (A-B) Expose the tooth extraction site. (C-D) Use a disposable syringe to separate the gingiva. (E-F) Use the tip of the micro-forceps to loosen left M1. (G-H) There is a gap between the two molars. (I-J) Remove teeth. (K-L) The extracted M1 tooth and the socket in the oral cavity for tooth extraction.

**Note:** Use small cotton balls that are only slightly moist and do not drip to pour or squeeze. If necessary, use dry cotton balls to remove any excess residual liquid.

Be extremely careful to prevent the liquid from flowing into the throat.

16. One hand pulls the left buccal mucosa of the mouse, and uses disposable syringe needles to separate the gingiva of left M1 (Figure 3C-D).

**Note:** Be gentle with the movements to avoid injury to blood vessels and causing excessive bleeding.

17. Close the tips of the bent micro-forceps to pass through the gap between the first and second molars on the lower jaw (Figure 3E-F). While rotating along the long axis of the instrument, forcefully close the tip of the micro-forceps to loosen left M1.

**Note:** Avoid applying too much force during the clamping process to prevent damaging the tooth crown. Loosen the crown of left M1 towards the buccal side firstly, and then loosen it towards the labial side.

18. Use the bent micro-forceps to firmly grasp the root bifurcation of left M1, apply force along the long axis of the tooth, and perform lateral movements in both the buccal and palatal directions (Figure 3I-J). During the shaking process, apply vertical force towards the occlusal surface to dislodge the tooth and complete the extraction.

**Note:** The force applied during clamping should not be too strong to avoid crushing the dental crown. Don’t bite into the gum to avoid causing bleeding of the soft tissues. When shaking, carefully feel the degree of looseness between the alveolar bone and the teeth. The shaking should be gradually intensified. When there is no obvious resistance, try to apply force vertically towards the occlusal surface.

19. Check whether the tooth root is intact, and whether there are any fragments of alveolar bone or large masses of soft tissue attached to it. Clean the surgical area with normal saline.
20. Following the above steps, remove the right M1 similarly.

**Note:** Each mouse needs to have all the instruments disinfected before the tooth extraction surgery.

Postoperative care

**Timing: [1-2 hours per mouse]**

21. After the surgery, gently remove the mouse from the fixed surgical plate. Drop a drop of normal saline on the tip of the mouse’s tongue to moisten its mouth. **Note:** The amount of normal saline dropped should not be excessive, as excessive liquid may enter the respiratory tract, causing the mouse to cough and suffocate.
22. Place the mouse on the heating pad and keep it in a prone position. The head should be tilted to one side to prevent liquid from entering the mouse’s respiratory tract.
23. Maintain this position until the mouse regains consciousness. Continuously observe whether the mouse has regained consciousness, whether there is breathing difficulty, and whether there is continuous bleeding in the surgical area.

**Note:** This process usually lasts for 1-2 hours. During this period, a drop of normal saline can be applied to the tip of the mouse’s tongue every half an hour.

24. Transfer the animals to clean cages and feed the mice in standard cages with a 12 h light/dark cycle and ad libitum access to food and water. Provide fresh and softened food.

Postoperative drug injection (Only For BRONJ)

**Timing: [2 weeks]**

25. After the surgery, continuous medication injections is required for 2 other weeks, with the prepared solution at a dose of 200 µg/kg of ZA, three times a week as before.

**Note:** Continuously monitor whether the mice show signs of swelling, bleeding, infection or pain. If any severe cases are found, euthanasia should be carried out immediately.

Tissue collection

**Timing: [15-20 minutes]**

26. After the mice complete 8 weeks of drug injection, euthanize the mouse using carbon dioxide (CO2) inhalation until they stopped moving. Use cervical dislocation to ensure the mice totally death.
27. The mandibles of the mice were collected and the soft tissues on the bones were scraped off.

**Note:** During the collection process, first cut open the skin and muscles on the cheek of the mice, exposing the bone surface. Then, cut off the mucosa and muscles attached to the inner side of the mandible from the mouth of the mice. Be careful to separate the condyle of the temporomandibular joint. Gently remove the soft tissues on the bone surface using scissors and forceps. Be careful not to damage the new bone tissue in the extraction socket.

28. Immerse the mandibles of the mice in 4% Paraformaldehyde solution at 4°C for 3 hours. Rinse 3 times with PBS for 5 minutes each. Afterwards, store the samples in PBS at 4°C.

Micro-CT analysis

**Timing: [1-2 hours per sample for micro-CT; 20-30 minutes per sample for morphometric analysis]**

29. Remove the mandibles from the PBS and place it in the micro-CT scanner with preset parameters. This protocol uses a 34mm diameter sample tube for scanning, with Voxel size 10.0 μm, FOV 35.36 mm, Image matrix 3400×3400×775, Slices 775, Voltage 55 kVp, Intensity 145 μA. Return the scanned samples in PBS for storage.
30. Reconstruct 3D images using the manufacturer’s software (DataViewer and CTvox). Lower and upper gray threshold values set to 130 and 255.

**Note:** The region of interest (ROI) is defined as the extraction socket of the first molar on the mandible.

Histology or immunofluorescence analysis

**Timing: [1 week]**

31. Decalcify in 10% EDTA (pH 7.4) for 5 days at 4 °C, changing the solution every day. Wash the samples 3 times using PBS for 10 minutes each.

**Note:** Use samples only scanned by micro-CT or new ones. Confirm that the mandible has been completely demineralized before proceeding with the subsequent steps. The method is to insert a disposable syringe needle into the thicker part of samples. If the needle can be inserted easily, decalcification is successfully achieved. Process samples as follows:

For histology

32. Process the samples by the automatic tissue processor treated with ethanol of different concentrations (70%, 80%, 95% and 100%) first and then xylene.
33. Insert the maxilla into the embedding mold and confirm that the sagittal plane is correct. Quickly inject 60 °C paraffin and gently press the specimen to prevent air bubbles. Cool at room temperature until complete solidification.

**Note:** The longitudinal axis of the molar row is usually the key observation plane for the sagittal section. Store at 4 °C for a short period for easy slicing.

34. Sectioning. The thickness of the section is 3 μm. The section is attached to a positively charged adhesive slide. This protocol uses H&E staining to evaluate the modeling effects of the model of BRONJ in mice.
35. Place and scan in the digital slide scanner.

### For immunofluorescence analysis

36. The samples are subjected to dehydration treatment with 20% sucrose and 2% polyvinylpyrrolidone in 4 °C for 24 hours. Then embed the samples 20% sucrose, 2% polyvinylpyrrolidone and 8% gelatin. Cut the samples into 80 μm thick sections using a low-profile blade on the cryostat and dry in air before being stored in a freezer in - 20 °C.

**Note:** Place embedded samples in a −20 °C freezer for freezing to facilitate faster solidification.

37. Perform permeabilization with Triton X-100, block with 5% horse serum, and conduct immunostaining with antibody solution. Add the nuclear marker DAPI to the secondary antibody solution for staining.
38. Place and scan in the spinning disk confocal microscope.
39. The images captured on the digital slide scanner are processed using OlyVIA. The images captured on the spinning disk confocal microscope are processed and reconstructed using Imaris and Imaris File Converter. Adobe Illustrator software is used for image processing and analysis.

## TROUBLESHOOTING

### Problem 1

Inadequate radiation injury (step “Animal preparation”, 5-6).

### Potential solution

Ensure sufficient anesthesia to avoid changed location of mouse mandible. Adjust the chamber lid to expose the mandible to radiation area.

### Problem 2

Inadequate anesthesia (step “Animal preparation”, 12).

### Potential solution

Inject the reagents strictly and slowly according to the dosage provided by the reagent manufacturer and the actual weight of the mice. Pay attention to the position of the mouse during the injection and the depth of the needle insertion. Do not make large movements that may cause bleeding in the abdominal cavity of the mouse.

Continuously monitor whether the reflexes of the mice exist (such as toe-pinch and righting reflex). If reflexes are observed, promptly replenish the dosage according to the instructions provided by the reagent manufacturer.

### Problem 3

Excessive bleeding during surgery (step “Tooth extraction”, 16-18).

### Potential solution

During the operation, be careful not to damage the soft tissues such as the gums, tongue, and mucosa with sharp metal instruments. The forceps used to open the mouth can be separated by several cotton balls and the mouse’s tongue to prevent mechanical damage. Be careful to insert the micro-forceps into the root bifurcation area gently and do not clamp the gums.

Prepare a certain number of dry cotton balls in advance. In case of accidental puncturing of blood vessels and bleeding during the surgery, these cotton balls can be used to promptly absorb the blood and immediately disinfect with povidone-iodine cotton balls. Pre-position several cotton balls in the oral cavity. If the operation is proficient, be careful not to affect the breathing of the mouse.

### Problem 4

Crown fracture (step “Tooth extraction”, 18).

### Potential solution

Avoid using sudden and excessive external force. Instead, apply controlled and gradual force. Find a stable fulcrum and you can appropriately generate force through the movements of your wrist and arm.

For Crown fracture, try to loosen the tooth root from the alveolar bone using a needle, and use a micro-forceps for a precise extraction. It is recommended to practice on a mouse cadaver before the actual surgery.

### Problem 5

Weight loss (step “Postoperative care”, 24).

### Potential solution

Provide appropriate soft food. If approved by the IACUC, consider using analgesic (buprenorphine 0.05-0.1 mg/kg SC). If the condition persists, humane endpoint should be applied to the mice.

### Problem 6

Death after radiation and tooth extraction (step “Preoperative management”, 6; “Postoperative care”, 24).

### Potential solution

Avoid excessive exposure to radiation and additional injury. Check the setting of irradiator to ensure proper radiation dose. Provide softened food after radiation and tooth extraction.

## EXPECTED OUTCOMES

Using this protocol, mice are expected to develop consistent and reproducible BRONJ within 8 weeks and ORNJ within 4 weeks. Compared with control mice (Figure 4A-B), BRONJ mice typically exhibit delayed or incomplete mucosal healing with partial bone exposure at the extraction site (Figure 4D-E). Micro-CT and histological analyses further demonstrate impaired bone regeneration in the extraction socket with the presence of necrotic bone tissue (Figure 5A). Immunolabeling reveals reduced endothelial cell abundance and decreased osteogenic activity within BRONJ lesions (Figure 5C). In ORNJ mice, histological analysis demonstrates endothelial swelling, microthrombus formation, increased osteoclast activity, and enlarged bone lacunae (Figure 5B), consistent with radiation-induced bone damage and impaired bone remodeling.

**Figure 4.**
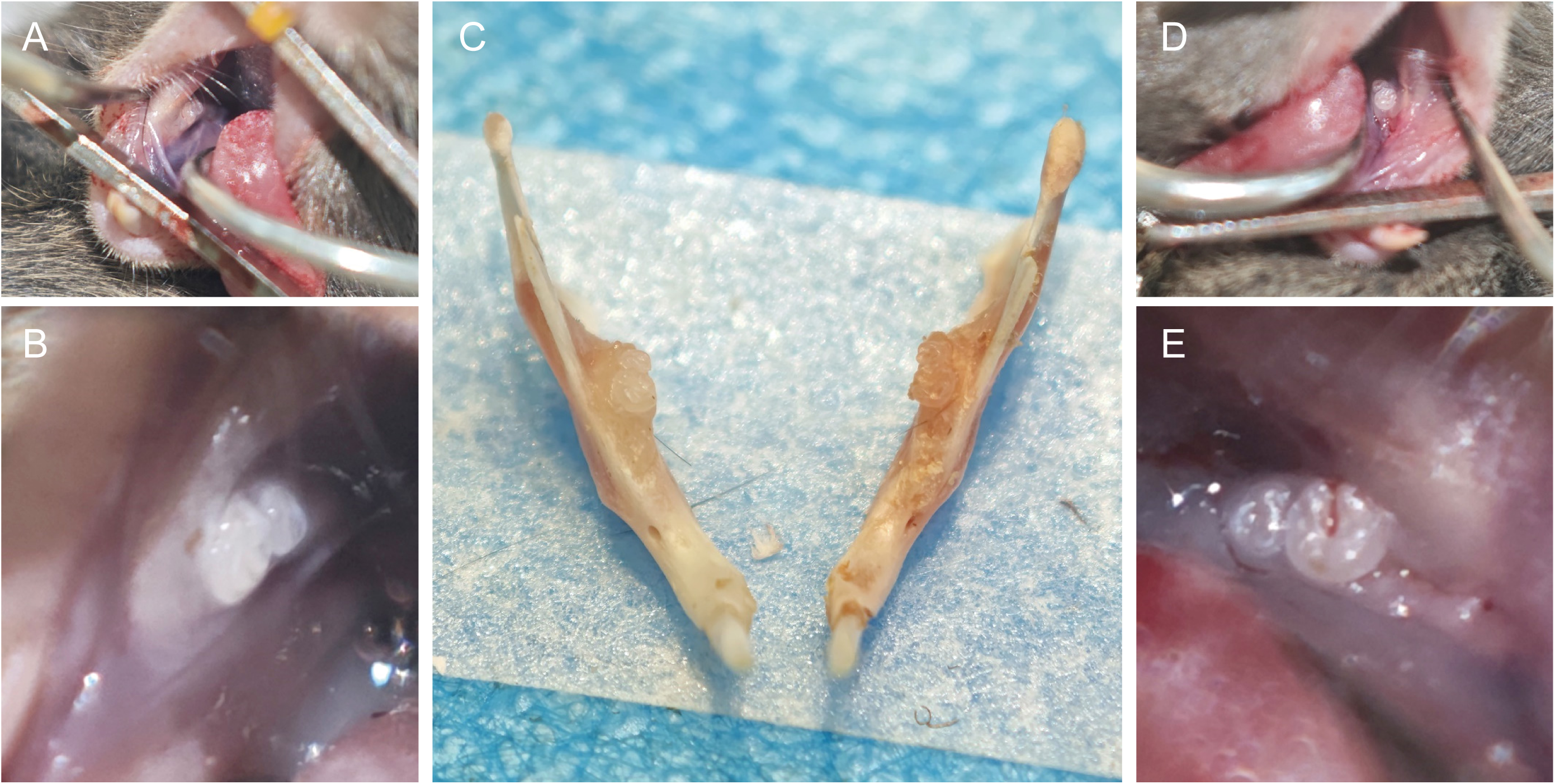
The comparison between samples of BRONJ and control. (A-B) Intraoral images of control. (C) Comparison of samples between the BRONJ group and the control group, with. (D-E) Intraoral images of BRONJ shows poor healing of the oral mucosa.

**Figure 5.**
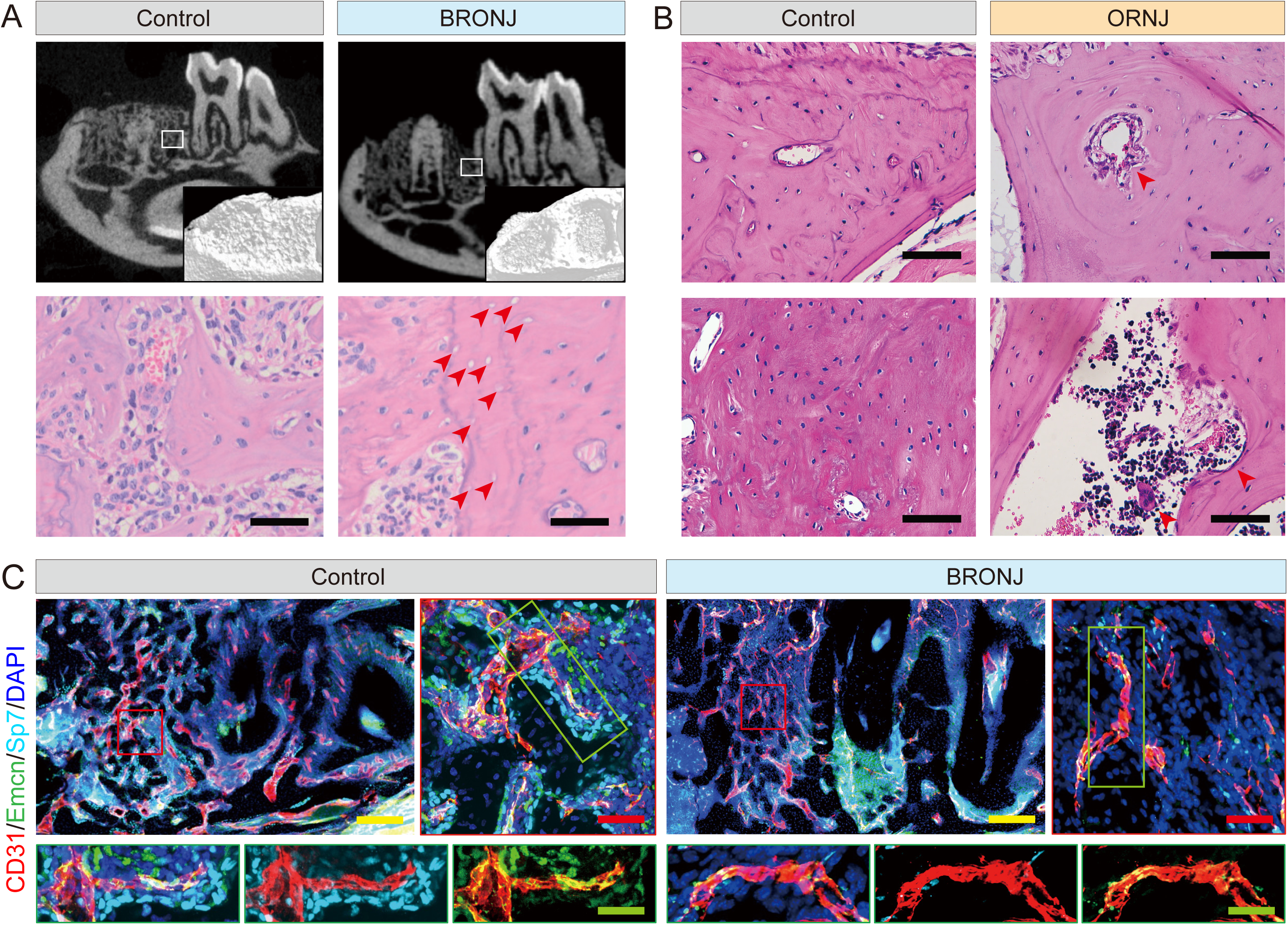
Analysis of the BRONJ and ORNJ compared with control. (A) Micro-CT shows lower bone density in BRONJ. Scale bars: 500 μm. H&E staining shows that pathological manifestations of bone lacuna. Scale bars: 50 μm. (B) The H&E staining of mandible sample sections shows that pathological manifestations of vasculature, osteoclast, and bone lacuna. Scale bars: 100 μm. (C) Immunolabeling in BRONJ shows reduced endothelial cell abundance and decreased osteogenic activity. Scale bars: 100 μm (yellow), 40 μm (red), 15 μm (green).

## LIMITATIONS

This protocol provides practical and reproducible steps for establishing mouse BRONJ and ORNJ models; however, several limitations should be considered. First, although detailed procedures for tooth extraction are provided, the surgery still requires advanced microsurgical skills. Inexperienced operators may encounter a higher failure rate, as improper manipulation can cause excessive trauma or even mandibular fracture, thereby compromising experimental outcomes. Second, this protocol is optimized for young adult mice. In older mice, particularly those aged over 50 weeks, age-related alterations in bone architecture may increase the risk of root fracture during molar extraction; notably, BRONJ lesions can still develop even when residual roots remain. Third, ZA administration alone does not fully recapitulate the broad clinical spectrum of medication-related osteonecrosis of the jaw. Fourth, a single high-dose irradiation regimen is employed to induce injury, which differs from the fractionated radiotherapy used in clinical practice. Although this approach may increase mortality due to extensive tissue damage, it enables more rapid induction of ORNJ. Despite these limitations, the protocol provides a robust and controllable platform for preclinical studies of jaw osteonecrosis.

## RESOURCE AVAILABILITY

### Lead contact

Further information and requests for resources and reagents should be directed to and will be fulfilled by the lead contact, Anjali P Kusumbe.

### Technical contact

Questions about the technical specifics of performing the protocol should be directed to the technical contact, Junyu Chen.

### Materials availability

This study did not generate new unique reagents.

### Data and code availability

This study did not generate or analyze datasets. This study did not report original code.

## ACKNOWLEDGMENTS

J. C. is supported by National Natural Science Foundation of China (Nos. 82422021, 82270961) and Sichuan Provincial Health Commission (24QNMP015). A. P. K. is supported by Ministry of Education (MOE) Singapore: Academic Research Funds (#024983-00001; #025277-00026), European Research Council (StG: metaNiche, 805201) and European Union’s Horizon 2020 (No 857524). A.C. is supported by the National Institutes of Health (NIH) under grant R01AG082681 (AC). M.V.R. acknowledges support by the National Institute on Aging (R01AG073349) and National Institute of Arthritis and Musculoskeletal and Skin Diseases (R01AR055655 and R01AR082460).

## AUTHOR CONTRIBUTIONS

Z.D., J.Z., and H.L. methodology, investigation, visualization, and writing. A.C. and M.V.R. reviewing, and editing. J.C. methodology, validation, and formal analysis. A.P.K. supervision, reviewing, and editing.

